# Preparatory Cortical Modulations for Stepping Tasks with Varying Postural Complexity

**DOI:** 10.1101/2025.07.28.667304

**Authors:** Ali Doroodchi

## Abstract

We examined whether preparatory cortical activity indexed by the contingent negative variation (CNV) scales with postural complexity during step initiation. Participants performed straight and diagonal stepping in a Warning-Go paradigm while EEG was recorded; CNV epochs spanned the 2s fore period and were summarized into eight 0.25-s bins for electrode and eLORETA source-level analyses using linear mixed-effects models (n=31).

Diagonal stepping produced greater early CNV negativity at the scalp (bin 1: C1, CP3, CP1, P1, FC4; bin 2: F1, F3, FC1, Fz, F2, F4, FC2, FCz), with no electrodes favoring straight stepping.

Source analysis showed stronger engagement for diagonal stepping in bins 1-3 (0-0.75 s) across fronto-parietal sensorimotor regions, including paracentral, transverse frontopolar, superior frontal (gyrus/sulcus), supramarginal, superior parietal, intraparietal, and precentral sulcus; no regions were greater for straight stepping. These effects concentrated in the early CNV suggest enhanced anticipatory selective attention and sensory up-weighting under higher postural demands, providing a richer state estimate for scaling anticipatory postural adjustments.

## Introduction

Before any movement is executed, the nervous system enters a preparatory phase (Ariani et al., 2022; Day & Bancroft, 2018). Sensory pathways selectively filter and modulate incoming inputs to amplify information most relevant to the impending task, while motor circuits establish a readiness state that organizes forthcoming commands (Ariani et al., 2022; Frigon & Rossignol, 2006). In this paper, we focus on these complementary processes, sensory processing and motor preparation, as they relate to postural control during step initiation.

## Stepping Initiation

Initiating a step requires maintaining dynamic balance while generating forward propulsion (Bancroft & Day, 2016; Day & Bancroft, 2018). To achieve this, the nervous system deploys anticipatory postural adjustments (APAs) that precede foot-off to counter the destabilizing effect of limb unloading by shifting body weight and create the initial propulsive impulse (Bancroft & Day, 2016; Farinelli et al., 2021). APAs rely on both motor planning and sensory integration (Aydin et al., 2022). Motor planning specifies the onset and magnitude of postural muscle activation (Ariani et al., 2022). Sensory integration weights inputs from variety of sensory modalities (Brown & Staines, 2015). The combined effect is a forward shift of the center of mass that remains within the changing base of support (Bancroft & Day, 2016). Together, these preparatory processes establish the initial conditions for a stable, efficient step (Bancroft & Day, 2016; Day & Bancroft, 2018).

## The contingent negative variation (CNV)

The contingent negative variation (CNV) is a slow cortical potential that reflects the allocation of neural resources for preparing an impending action, appearing as a gradual negative shift on scalp electroencephalograph (EEG) (Fattapposta et al., 2024). In paradigms assessing the CNV, an initial warning cue establishes expectancy and gives information about an upcoming task, and a subsequent imperative cue signals movement execution (Fattapposta et al., 2024; A. W. K. Gaillard, 1976). The CNV comprises an early orientation wave (O-wave) that modulates sensory processing to receive task-relevant information into the brain, followed by a later phase proximal to the imperative cue that indexes motor planning and the organization of the forthcoming movement (A. Gaillard, 1977; Rockstroh et al., 1993). Together, these phases capture the transition from anticipatory sensory gating to motor preparation to ready the individual for an expected movement.

During the early CNV (orientation wave), anticipatory selective attention from prefrontal control regions biases thalamocortical gating, sharpening sensory filtering and enhancing activity in task-relevant sensory cortices to improve the fidelity of incoming information (Dale et al., 2008). As the imperative approaches, the later CNV reflects motor preparation: premotor, supplementary motor, and primary motor areas ramp to specify the timing and scaling of forthcoming commands, while fronto-parietal circuits integrate relevant sensory inputs into the motor plan (Alyahyay et al., 2023; Brunia, 1993). Together, these mechanisms transition the system from enhanced sensory intake to organized action.

Applied to step initiation, these mechanisms serve two complementary roles. Anticipatory selective attention enhances plantar cutaneous inputs that track center of pressure (CoP) and inform center of mass (CoM) estimation, yielding a higher-fidelity state estimate on which APAs are computed (Ariani et al., 2022; Kavounoudias et al., 1998). When a task makes CoM monitoring more critical, this sensory heightening may be amplified further compared to a task where CoM monitoring is less critical (Mouchnino et al., 2015). The later CNV indexes allocation of neural resources for planning the APAs that unload the limb and generate forward propulsion (Bent et al., 2002). Tasks demanding more complex APA patterns may therefore recruit greater preparatory resources, producing a larger CNV (more negative) than tasks with simpler APAs (Delse et al., 1972; Kononowicz & Penney, 2016).

## Electroencephalography (EEG)

Electroencephalography (EEG) measures the electrical dipoles generated from pyramidal neurons collective activation (Gable et al., 2022). If a sufficient number of pyramidal neurons in close proximity are activated simultaneously, their collective dipole can propagate through brain’s tissues via volume conduction and be measured via EEG electrodes placed on the scalp (Gable et al., 2022; Hádinger et al., 2022). Cortical activity’s source cannot be estimated from the surface EEG electrodes because the dipoles created from different neurons interact with each other, and the individual dipoles propagation are tempered as they go through different tissues such as the brain, meninges, and the skull (Gable et al., 2022). Source localization algorithms like exact Low Resolution Electromagnetic Tomography (eLORETA) aim to estimate the most probable location of cortical activity that may give rise to the recorded activity at the electrode level, by considering the attenuation imposed by different head tissues and the possible ways the dipoles may interact with each other (Faes et al., 2021; Jatoi et al., 2014). Therefore, eLORETA analysis may be used to explore CNV’s cortical origins more exhaustively than what may be interpretable from the data from EEG electrodes located on the scalp (Faes et al., 2021; Jatoi et al., 2014).

## Research Direction

Although the kinematics for different stepping movements and their proceeding APAs have been explored before, it remains unclear how postural complexity may influence neuronal activity for stepping preparation’s motor planning and sensory modulation. We aimed to investigate whether CNV patterns differs with postural complexity, hypothesizing that greater postural complexity (e.g., in diagonal stepping) would create a larger (more negative) CNV.

## Methods

### Tasks Selection & Rationale

EEG recording was done while participants performed two stepping tasks: straight stepping and diagonal stepping. These stepping tasks were selected based on established differences in their APA kinematics (Bancroft & Day, 2016; Lyon & Day, 1997). Diagonal stepping requires a more pronounced asymmetrical CoP shift to meet the greater balance perturbations of the diagonal step vector, demanding greater postural control compared to straight stepping for safe and efficient execution (Bancroft & Day, 2016; Lyon & Day, 1997). Consequently, differences in motor planning processes across these tasks may reflect increased sensorimotor integration and preparatory neural activity as postural demands intensify. Additionally, since directional stepping is frequently performed in daily activities, the findings from this study can be readily generalized to real-world scenarios (Mille et al., 2013; Yates et al., 2023).

#### Task Instructions

Participants were instructed to adopt a natural and comfortable stance. Participants were asked to step forward to a length that felt natural for initiating walking. For diagonal stepping trials, the step was performed approximately 45 degrees lateral from the forward direction. All Stepping trials were done with the right foot as the stepping foot, and the left foot as the stance foot.

Participants were instructed to maintain stepping characteristics such as step length, speed, and angle, across trials as consistent as possible across trials. To minimize eye and head movement artifacts, a fixation point was placed at eye level on the wall directly in front of the participants, who were instructed to maintain their gaze on this point and keep head movements to a minimum.

Participants took a step, held the stepping foot in its position for 3 seconds, and then returned to their original stance. This brief hold was introduced to ensure the participants transferred their weight to the stepping foot and performed the step fully. After returning to their original stance, the participants stood stationary for 7 seconds prior to hearing instructions regarding the next step. Limiting the stepping to one step was due to equipment constraints, as the EEG cables were not long enough to safely allow for more than one step. However, there are no differences in APAs beyond the first step, so taking a single step was sufficient for investigating the neural activations associated with the different APAs’ postural demands (Farinelli et al., 2021). All steps were performed with the right leg and the left leg was the stance leg. The left leg was free to move to accommodate the step, but it never fully left the ground. Multiple practice steps were performed, and recording started once the participant understood the paradigm. If the participants deviated from the instructions, they were reminded of the correct procedures as appropriate.

#### Stepping Trial Quantities

The stepping trials consisted of 40 trials per stepping task, totaling 80 steps. The tasks were performed in 2 sets of 40, with the option of a 5-minute break between each set. The order of the stepping tasks (straight vs. diagonal) in each set was randomized until the maximum number of trails for either of the stepping tasks in the set (i.e., 40) were reached. After the maximum number of trials for a stepping condition was reached, the remainder of the stepping tasks in the set were switched to the unfinished task. This randomization was done live in the session using a Python script. The entire stepping paradigm lasted about 45 minutes, with each set of 40 steps taking approximately 8 minutes. The entire experiment, including EEG setup, familiarization with the protocol, and the consent process, took between 45-60 minutes.

### Sample

A total of 40 participants were recruited from the local population who met the following inclusion and exclusion criteria. Participants were included if they were between the ages of 18-40, and able to fluently communicate in English. Participants were excluded from this project if they had any current lower extremity musculoskeletal injuries/conditions (previous musculoskeletal injuries were accepted as long as recovered), lower extremity sensory injuries/conditions, neurological conditions that might affect gait or attention (such as Attention deficit hyperactivity disorder, Parkinson’s disease, stroke, etc.) circulatory problems in the lower extremity (e.g., Deep-vein thrombosis, impaired circulation), electronically active implants (e.g., pacemakers, cardiovascular defibrillators) or metallic implants in the foot.

Participants were recruited through the OurBrainsCan Registry and through a mass recruitment email from Western University’s registry. The University of Western Ontario’s research board committee approved this study. After enrolment, we were unable to book 5 participants’ data collection for this project. Of the remaining 35, 3 participants’ data were eliminated due to equipment technical difficulties, and 1 was eliminated due to deviations from the study protocol. 31 participants data are included in this project. The included sample size’s average age was 23.71 years (SD = 4.95), with 17 males and 14 females.

### Cues

Auditory cues were played through speakers. The auditory cues included a ‘*Warning’* cue and a ‘*Go’* cue. The warning cue informed participants about the upcoming stepping task (either straight or diagonal). The warning auditory cues were “One”, associated with forward stepping, and “Two”, associate with diagonal stepping. Two seconds after the termination of the Warning cue, a 440 Hz beep sound was played as the Go signal, instructing the participants to execute the stepping task. Participants were instructed to be prepared for the step after hearing the warning cue (i.e., “One” or “Two”) but not to initiate the step until they hear the imperative cue (the 440 Hz beep). The participants were instructed to maintain their position after stepping until they hear the return cue. An auditory cue “Return” was played 3s after the termination of the Go cue to instruct the participants to return to their original stance.

### EEG

The Biosemi Active 2 64-channel EEG system (Netherlands) was used with a sampling rate set to 1024 Hz. The 10-20 international montage system was used, and the electrode impedance was reduced to less than 10 microohms. Data collection was performed in an electromagnetic and acoustic shielded room.

### EEG Data Pre-Processing

The EEG file was imported into Python (version 2024.12.3) and analyzed using the mne library (version 1.7) (Larson et al., 2024). The following operations were done in the order described below. Bad channels were removed, and their data was interpolated based on the remaining good channels. All electrodes were referenced to the average signal across all electrodes in their respective file. Notch filters at 60 Hz and its harmonics (60, 120, 180, 240, 300, and 360 Hz) were applied using a finite impulse response (FIR) filtering method to remove line noise. After notch filters’ application, Independent Component Analysis (ICA) with the fastICA algorithm was used for artifact removal to the greatest extent possible. Following artifact removal with ICAs, the data was filtered with a high-pass frequency of 0.1 Hz and a low-pass frequency of 40 Hz.

#### Epoching

The data from warning stimulus presentation to 2 seconds after its presentation was averaged to obtain the CNV epochs. 0.1s prior to the Warning stimulus presentation until its onset served as each epoch’s base line period. The average value in each baseline period was subtracted from its respective epoch.

##### Epoch Rejection Criteria

First, epochs were also excluded if participants performed the task incorrectly or if there were deviations from the experimental protocol. This included instances where movement timing was inconsistent with task instructions (e.g., stepping on the warning cue), performing the wrong movement (e.g., stepping diagonally when instructed to step straight) or when other factors interfered (e.g., participant talking during the trials, phone ringing) with the expected execution of the task. By removing these epochs, it was ensured that only trials reflecting proper task execution and adherence to the study design were included in the final analysis, enhancing the reliability of the results. Removing these bad epochs resulted in an unequal number of good epochs across participants as the epoch’s exclusion criteria did not affect all participants similarly.

To increase signal to noise ratio in the good epochs remaining, 15% of the nosiest epochs were also removed. To identify and remove noisy epochs, an amplitude-based thresholding method was applied (Mumtaz et al., 2019). Epochs containing signals at any of the electrodes that exceeded a specific threshold were marked as noisy. This threshold was not fixed but was iteratively adjusted to ensure the removal of 15% of the total epochs, which were the noisiest segments of each participant’s data. By dynamically setting this threshold, only the most contaminated epochs were excluded, preserving as much clean data as possible in a participant-specific manner. If an electrode contributed to the removal of a significant portion of the epochs and its data deemed noisy with visual inspection, that electrode was removed, interpolated, and ICAs and epoch removal were then repeated with the removed electrode’s interpolated data. Clean epochs’ weighted averaging provided each condition’s evoked response. Figure 1. Summarizes the pipeline prior to statistical analysis.

**Figure 1:**
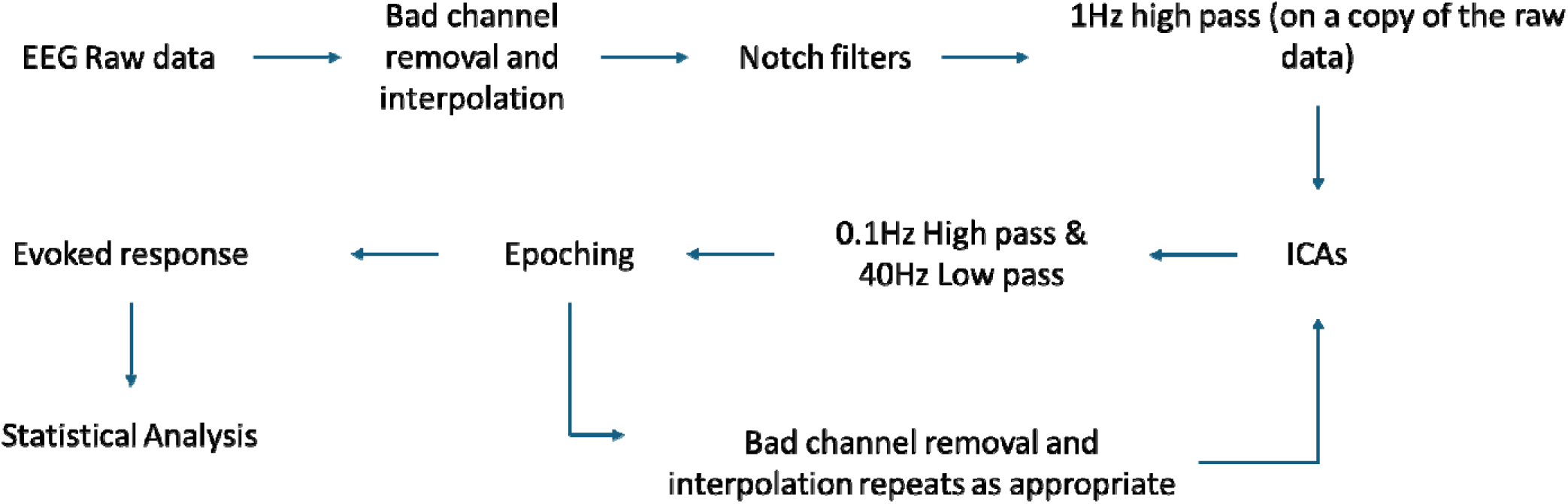
EEG Pre-processing Steps

### Source Localization

Good epochs and their evoked data were imported for source localization analysis. The noise covariance was created based on the epoch’s base line period (0.1s until the Warning cue). A forward solution was created based on the Free Surfer average (fsaverage) brain’s source spaces, transformation file and boundary element method (Wu et al., 2018). Based on the forward solution, an inverse operator was created and the its inverse solution was applied to the evoked data using the eLORETA weighted minimum norm inverse solution (Pascual-Marqui, 2007). To manage the tradeoff between data fidelity and noise sensitivity in the inverse solution, the square of the regularization parameter λ (lambda) was used, i.e.:

*Equation 1: was used as the noise sensitivity’s regularization parameter. SNR: Signal to Noise Ratio*

Based on the previous literature, a Signal to Noise Ratio (SNR) of 3 was deemed appropriate for the epoched data in such academic setting (Barzegaran et al., 2019; Beltrachini et al., 2021). The ‘*Automatic PARCellation, Destrieux Atlas, 2009, surface-based’*, was used to extract the mean regional activity for each sample frame across the evoked time window (Pascual-Marqui, 2007).

The following process, following literature’s guidelines, was used to parse out relevant cortical regions with sufficient cortical activation (Barzegaran et al., 2019; Beltrachini et al., 2021). First, a threshold was calculated by determining 80% of the activation amplitude in the region maximally activated for each component for each participant. Regions that met the 80% threshold in at least 50% of the participants were kept for further analysis. The activity in the preserved regions were compared between stepping conditions using the linear mixed effects model.

#### Statistical Analysis

After the preprocessing steps above, participants’ evoked data and source localization data were exported to .csv files and imported to R Studio for statistical analysis. To evaluate condition related differences in the evoked data and the source localization data, a linear mixed-effects model was used. The model included Condition as a fixed effect and a random intercept for anonymized participants’ ID to account for within-subject variability due to repeated measures. This approach allowed for modeling individual baseline differences while properly accounting for the correlation between repeated measurements from the same participant. The linear mixed-effects models were tested for linearity, normal distribution, homoscedasticity, and points overly influencing the regression coefficients by examining their diagnostic plots. Multicollinearity was checked by the generalized variance inflation factor and observation’s independence by the Durbin-Watson test. No discrepancies to the model’s assumptions arose from examining these diagnostic plots and tests.

### CNV Evoked Potential Analysis

For CNV’s statistical analysis, each electrode and cortical region mean activity for 0.25s time windows were calculated (e.g., mean activity at the Cz electrode, or Paracentral Gyrus & Sulcus in the right hemisphere at 0s - 0.25s, 0.25s - 0.5s…). Averaging the activity into 8-time bins allowed for separate analysis of motor preparation’s different stages while also improving the signal to noise ratio. Then the linear mixed effects were used to explore the differences in motor preparation between the Stepping tasks.

## Results

### Electrode Level Differences

The O-wave is visible on the CNV traces as indicated by the negative deflection following the auditory evoked potential, approximately occurring at 0.3s to 1.3s. Electrode level analysis highlighted the mean activity at electrodes C1, CP3, CP1, P1, and FC4 to be more negative at bin 1 (0s – 0.25s), and electrodes F1, F3, FC1, Fz, F2, F4, FC2, and FCz to be more negative at bin 2 (0.25s – 0.5s). Figure 2 shows the location of these electrodes, with their assigned colour highlighting the bin in which they showed more negativity. Panel B on Figure 1 shows the time frames these electrodes showed significantly more negativity, with the line’s colours being matched to the electrode’s colours in Figure 2. Table 1 shows the statistical information for these electrodes, with the column colour matching Figure 2 Panel B and Figure 3. None of the electrodes showed significantly more negative activity in the straight stepping condition, compared to the diagonal stepping condition for CNV electrode level analysis.

**Figure 2:**
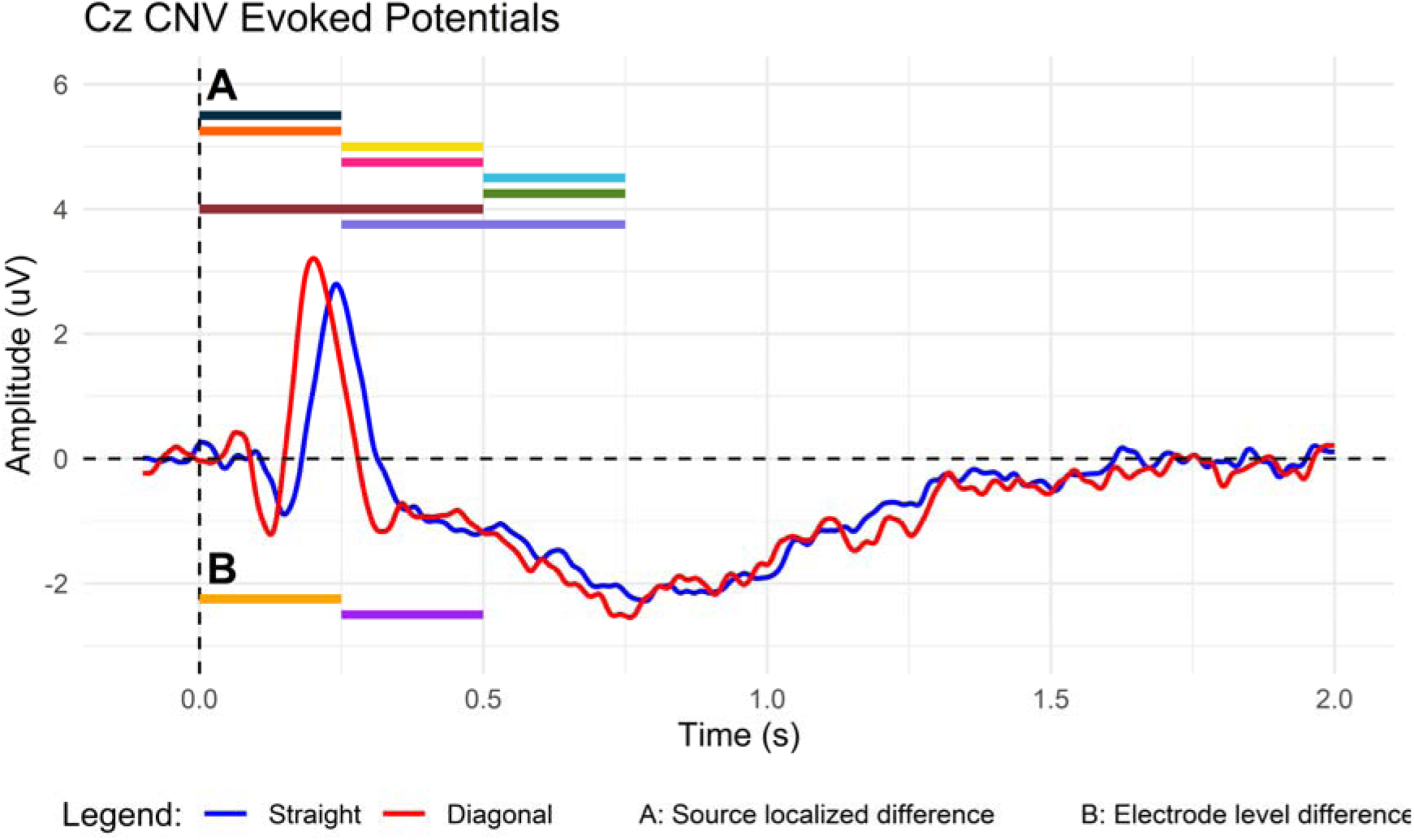
Blue and red traces highlight the CNV evoked potential from the Cz electrode for straight and diagonal stepping respectively. A: Regions that are significantly more active revealed by source localization analysis, placed on time bins were more negative.

**Figure 3:**
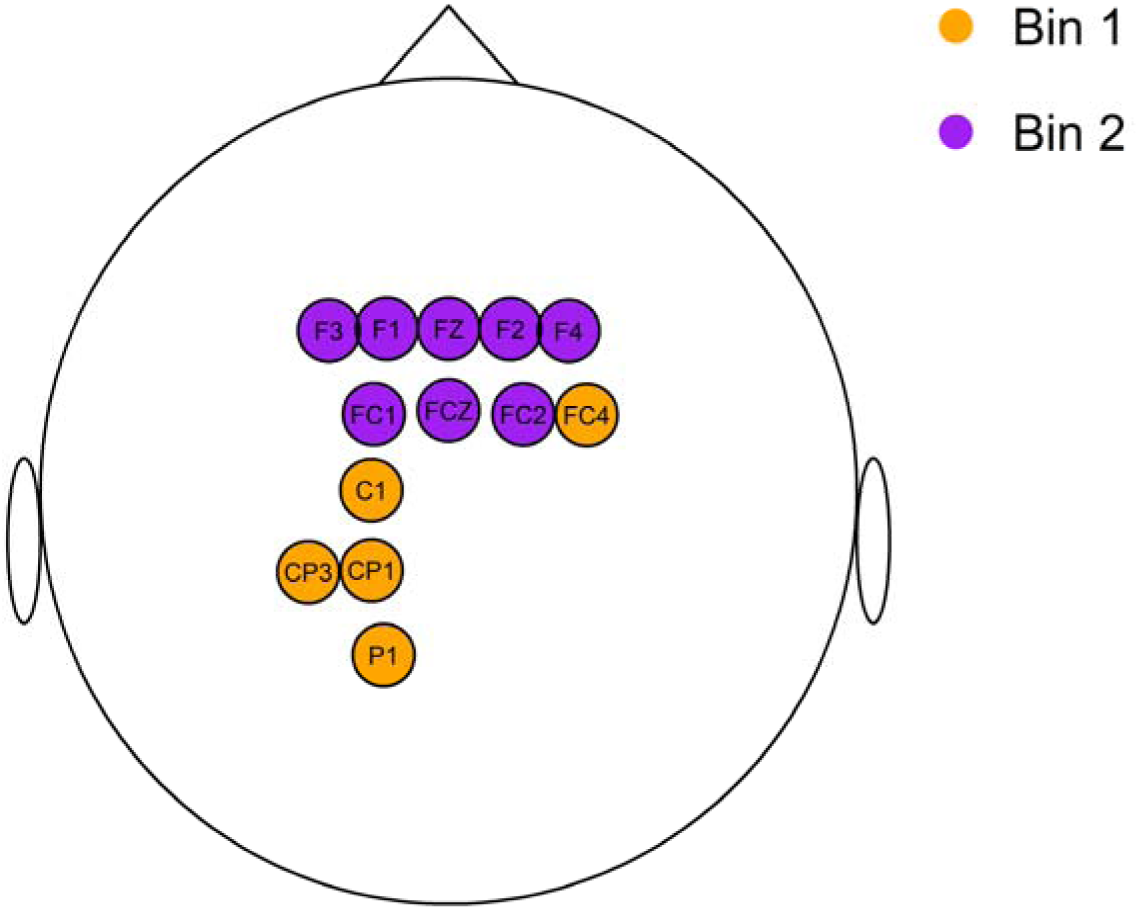
Electrodes that showed significantly different electrodes in CNV analysis. The colours match to the “Colour” column in Table 6.

**Table 1:**
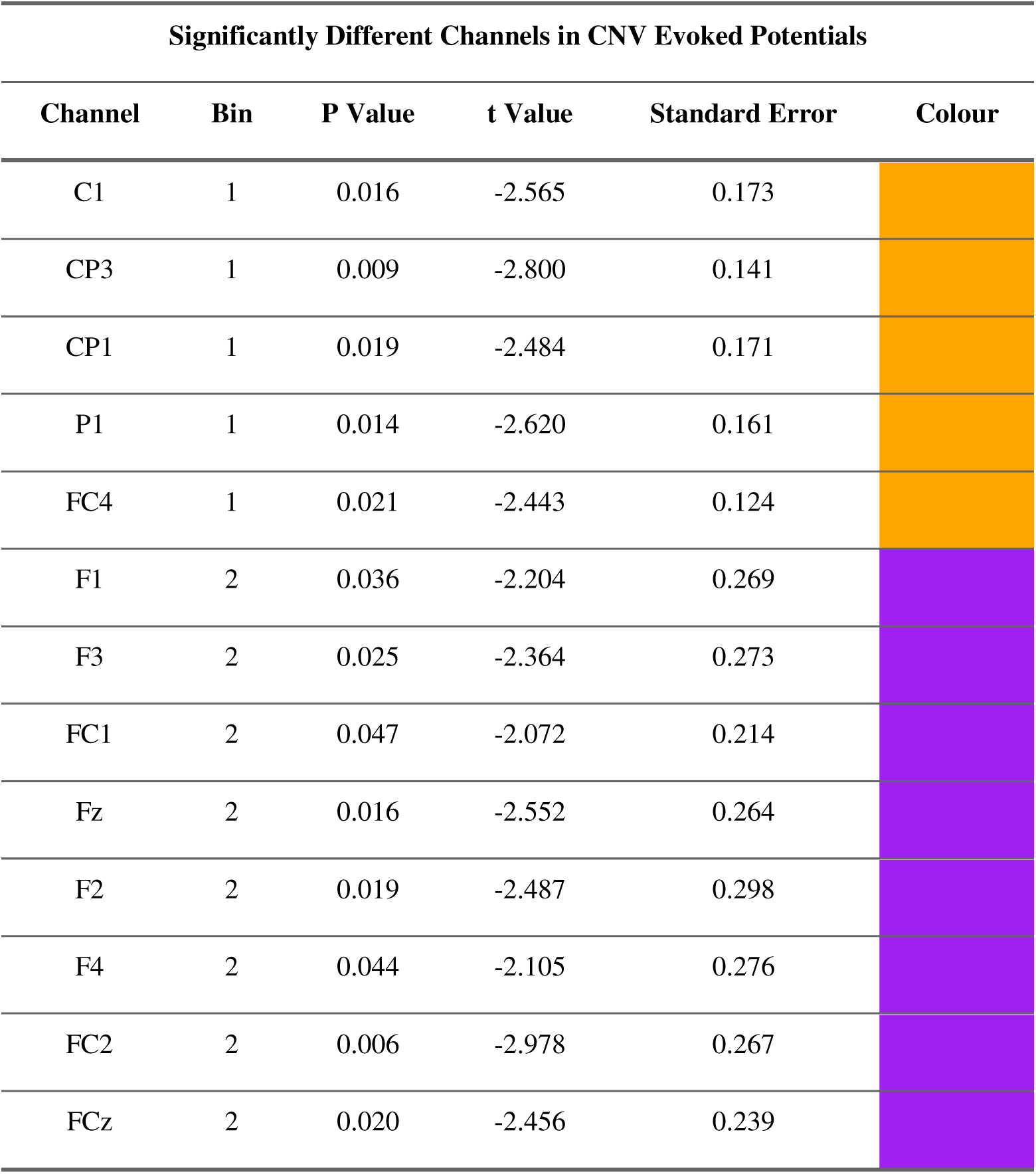
Statistical information for significantly different electrodes. The column “Colour” matches the electrode colours in Figure 13.

| Significantly Different Channels in CNV Evoked Potentials |  |  |  |  |  |
| --- | --- | --- | --- | --- | --- |
| Channel | Bin | P Value | t Value | Standard Error | Colour |
| C1 | 1 | 0.016 | -2.565 | 0.173 |  |
| CP3 | 1 | 0.009 | -2.800 | 0.141 |  |
| CP1 | 1 | 0.019 | -2.484 | 0.171 |  |
| P1 | 1 | 0.014 | -2.620 | 0.161 |  |
| FC4 | 1 | 0.021 | -2.443 | 0.124 |  |
| F1 | 2 | 0.036 | -2.204 | 0.269 |  |
| F3 | 2 | 0.025 | -2.364 | 0.273 |  |
| FC1 | 2 | 0.047 | -2.072 | 0.214 |  |
| Fz | 2 | 0.016 | -2.552 | 0.264 |  |
| F2 | 2 | 0.019 | -2.487 | 0.298 |  |
| F4 | 2 | 0.044 | -2.105 | 0.276 |  |
| FC2 | 2 | 0.006 | -2.978 | 0.267 |  |
| FCz | 2 | 0.020 | -2.456 | 0.239 |  |

Source Localization Analysis for the 8 CNV time bins highlighted the following cortical regions to be significantly more active in the diagonal stepping condition compared to straight stepping in bins 1-3 (0s - 0.75s): Paracentral Gyrus & Sulcus, Transverse Frontopolar Gyrus, Superior Frontal Gyrus, Supramarginal Gyrus, Superior Parietal Lobule, Superior Frontal Sulcus, Interparietal Sulcus, and Precentral Sulcus. Figure 2 shows the location of these cortical regions, with their assigned colour highlighting the bin in which they showed more negativity. The region of their significantly more activity is shown on figure 2 panel A, with the same colour assignment to figure 4. Table 2 shows the statistical information for these electrodes. No cortical region showed more activity in the straight stepping condition compared to the diagonal stepping condition during CNV source localization analysis.

**Figure 4:**
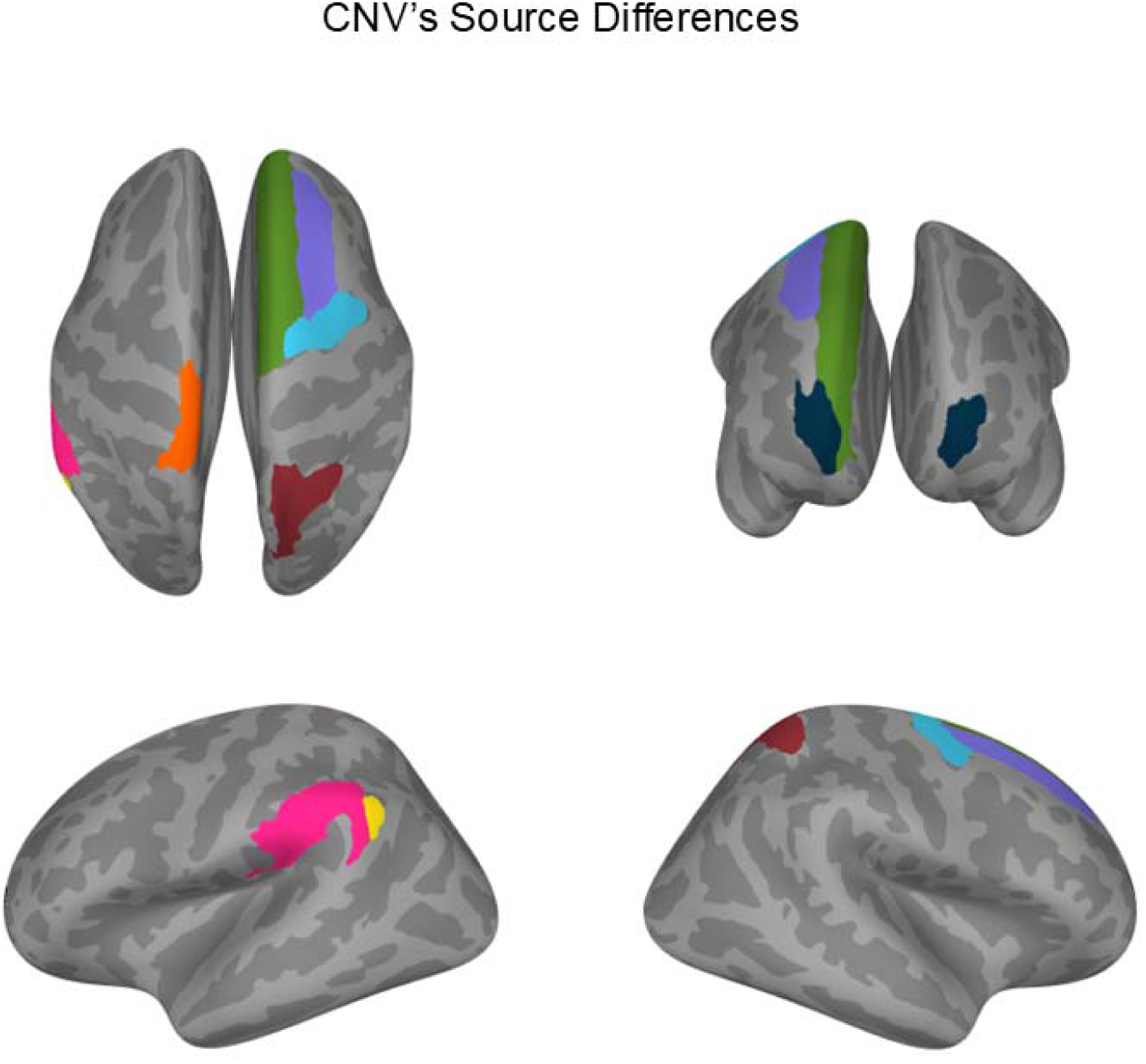
Significantly different regions revealed from CNV’s source localization analysis. The colours match to the “Colour” column in Table 8

**Table 2:**
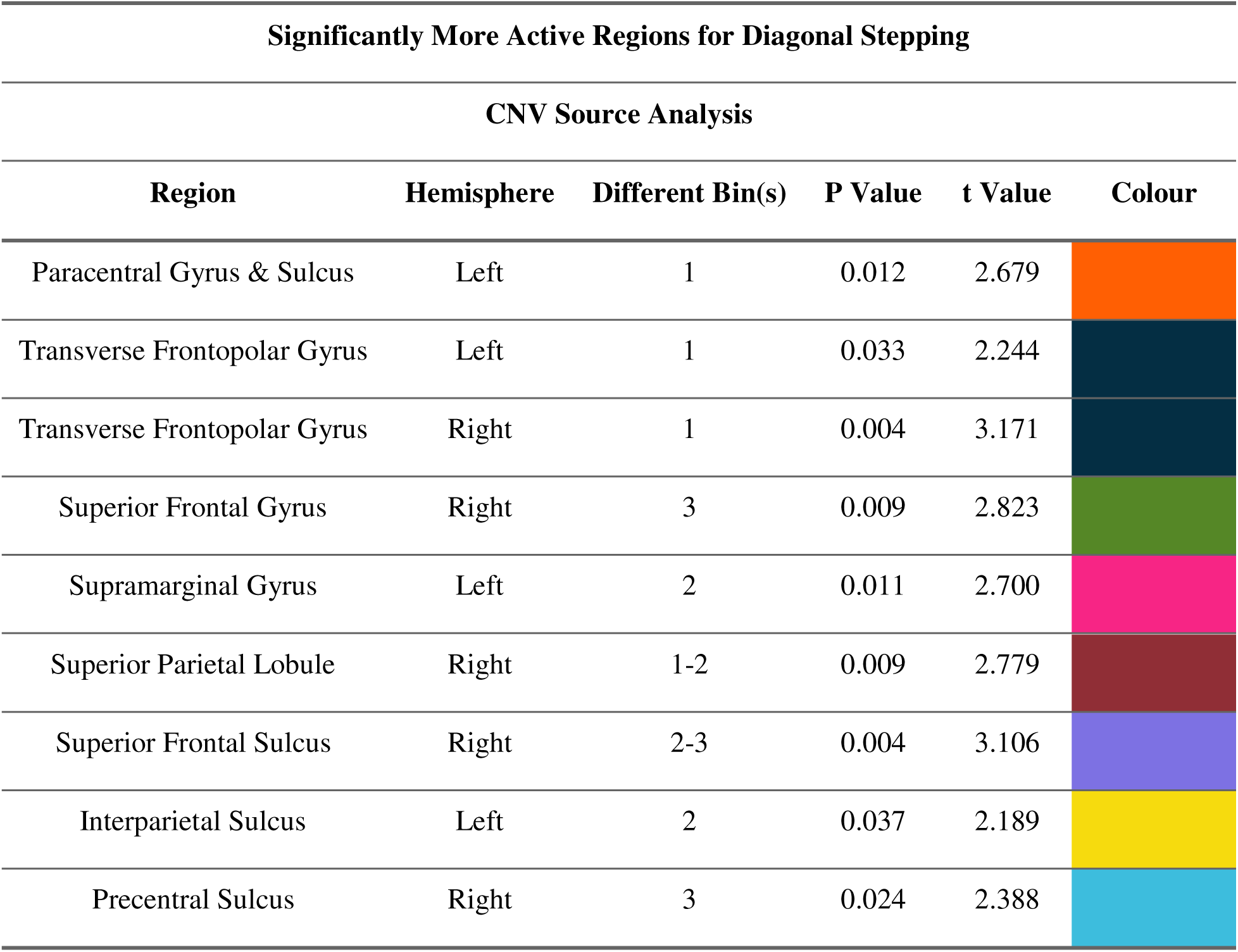
Statistical information for significantly more active cortical regions. The column “Colour” matches the electrode colours in Figure 13.

### Source Localization

Source Localization Analysis for the 8 CNV time bins highlighted the following cortical regions to be significantly more active in the diagonal stepping condition compared to straight stepping in bins 1-3 (0s - 0.75s): Paracentral Gyrus & Sulcus, Transverse Frontopolar Gyrus, Superior Frontal Gyrus, Supramarginal Gyrus, Superior Parietal Lobule, Superior Frontal Sulcus, Interparietal Sulcus, and Precentral Sulcus. Figure 4 shows the location of these cortical regions, with their assigned colour highlighting the bin in which they showed more negativity. The region of their significant more activity is shown on Figure 2 panel A, with the same colour assignment to Figure 4. Table 1 shows the statistical information for these electrodes. No cortical region showed more activity in the straight stepping condition compared to the diagonal stepping condition during CNV source localization analysis.

## Discussion

Following the warning cue, the waveform showed a clear auditory-evoked response, consistent with early processing of the cue. Immediately afterward, the signal transitioned into a sustained, gradual negativity that persisted through the fore period, the CNV. This sequence, an initial auditory response followed by a slow negative shift—frames our interpretation of the data. The early portion aligns with orienting and sensory gating, and the later portion reflects motor preparation for the upcoming step. In the discussion below, we assess how task-related postural complexity modulates each phase.

### Cortical differences

In the early CNV phase, both scalp and source analyses diverged by condition: diagonal stepping produced stronger negativity at the electrodes and greater engagement of frontal control, somatosensory, and parietal regions than straight stepping. The concentration of these effects in the early CNV points to differences in the orientation wave, consistent with heightened anticipatory selective attention to task-relevant inputs.

### Frontal Areas

Greater early frontal activity suggests stronger top-down attentional control (Haegens et al., 2011; Liu et al., 2016). This would increase gain and selectivity in task-relevant sensory cortices, raising the signal-to-noise ratio for inputs needed to initiate a step, especially plantar cutaneous and proprioceptive cues that inform CoP/CoM estimates (Mergner & Rosemeier, 1998; Shafiul Hasan et al., 2020). In the early CNV (orientation wave), this enhancement would support more precise state estimation for planning APAs.

Frontal regions can also influence sensory gating via corticothalamic control (Carmona et al., 2023; Crandall et al., 2015). Anatomically, the prefrontal cortex sends strong projections to the thalamic reticular nucleus (TRN), an inhibitory sheath that regulates thalamocortical transmission, and recent work shows region-specific cortical control over TRN circuitry (Hádinger et al., 2022). Functionally, TRN activity tracks attentional gating, while the pulvinar coordinates information flow and inter-areal synchrony according to attentional demands; more broadly, modern accounts place the thalamus as a hub that shifts and sustains cortical interactions during cognitive control (Alonan & Brown, 2002; Hádinger et al., 2022; Sherman, 2017). Together, these pathways provide plausible routes by which increased frontal activity in the early CNV could bias thalamocortical relay and selectively up-weight afferents most relevant for the impending step.

### Sensory Areas

Prefrontal control can bias thalamic relay via the thalamic reticular nucleus. This bias prioritizes task-relevant afferents before they reach cortex. Motor circuits, including premotor areas and M1, also set the state of primary somatosensory cortex (S1) (Ariani et al., 2022; Sherman, 2017). They establish sensorimotor expectations and suppress distracting inputs (Gómez et al., 2021). Together, these influences tune S1 toward the sensory consequences needed to plan anticipatory postural adjustments. The result is higher-fidelity CoP and CoM estimates to guide the upcoming step.

### Parietal Areas

Early engagement of the parietal cortex likely reformats sensory streams into action-ready codes (Andersen et al., 1997). Posterior parietal areas perform sensorimotor transformations, converting inputs represented in gaze- or skin-centered coordinates into body- and effector-centered frames useful for movement planning (Acimovic, 2008; Andersen et al., 1997). This includes tactile “remapping” that integrates proprioception and posture to place skin signals in external space, and gaze-centered visual updating that keeps spatial representations aligned as the eyes move (Akrami et al., 2018; Chao et al., 2015). In parallel, parietal circuits combine visual, proprioceptive, and vestibular cues to build a unified estimate of body state and orientation (Kaulmann et al., 2017; Medendorp et al., 2003). These computations make downstream motor planning more efficient by providing location, posture, and motion in coordinates that match the effectors (Mulliken et al., 2008).

### Implications for Diagonal stepping

Diagonal stepping imposes higher postural complexity, so APAs must be scaled and timed with tighter margins (Berchicci et al., 2020; Dingwell et al., 2023). Early frontal engagement sharpens sensory filtering, prioritizing step-relevant tactile, proprioceptive, and visual vestibular inputs while suppressing distractors (Aydin et al., 2022; Duque et al., 2013). This selective filtering yields a higher-fidelity body schema, more accurate CoP/CoM estimates and limb-load status on which APA commands can be computed (Zettel et al., 2002). Heightened S1 activity further increases the precision and reliability of these inputs (Borich et al., 2015). In parallel, parietal circuits transform and integrate them into leg- and body-centered, ground-fixed representations that are directly usable for foot placement and trajectory planning (Mulliken et al., 2008). Together, frontal filtering, S1 gain, and parietal re-coding provide the richer state estimate needed to scale APAs more intricately in the diagonal condition, accounting for the stronger early engagement observed in this task.

## Conclusion

Together, our results indicate that the nervous system flexibly scales preparatory cortical activity with postural complexity. When stepping demands more intricate control (as in the diagonal condition) early engagement increases across frontal, primary somatosensory, and parietal cortices. Heightened frontal control sharpens the selection of task-relevant afferents, S1 gain improves the precision of CoP/CoM estimates, and parietal transformations recode these inputs into action-centered coordinates (Edwards et al., 2019; Lyon & Day, 1997). This increased neuronal allocation yields a higher-fidelity body schema, ensuring that anticipatory postural adjustments are scaled and timed appropriately for a stable step (Andersen et al., 1997; Tommerdahl et al., 2010). The CNV thus provides a systematic index of this adaptive coupling between sensory modulation and motor preparation under heightened postural demands.

## Notes

### Competing Interest Statement

The authors have declared no competing interest.

